# Joint tensor modeling of single cell 3D genome and epigenetic data with Muscle

**DOI:** 10.1101/2023.01.27.525871

**Authors:** Kwangmoon Park, Sündüz Keleş

## Abstract

Emerging single cell technologies that simultaneously capture long-range interactions of genomic loci together with their DNA methylation levels are advancing our understanding of three-dimensional genome structure and its interplay with the epigenome at the single cell level. While methods to analyze data from single cell high throughput chromatin conformation capture (scHi-C) experiments are maturing, methods that can jointly analyze multiple single cell modalities with scHi-C data are lacking. Here, we introduce Muscle, a semi-nonnegative joint decomposition of **Mu**ltiple **s**ingle **c**el**l** t**e**nsors, to jointly analyze 3D conformation and DNA methylation data at the single cell level. Muscle takes advantage of the inherent tensor structure of the scHi-C data, and integrates this modality with DNA methylation. We developed an alternating least squares algorithm for estimating Muscle parameters and established its optimality properties. Parameters estimated by Muscle directly align with the key components of the downstream analysis of scHi-C data in a cell type specific manner. Evaluations with data-driven experiments and simulations demonstrate the advantages of the joint modeling framework of Muscle over single modality modeling or a baseline multi modality modeling for cell type delineation and elucidating associations between modalities. Muscle is publicly available at https://github.com/keleslab/muscle.

## 1 Introduction

Interactions between distal genomic regions (i.e., loci) that become in close proximity of each other through chromatin loops and topologically associated domains (TADs) are key elements of gene regulatory mechanisms. High-throughput Chromatin Conformation Capture (Hi-C) sequencing technology (Lieberman-Aiden et al., 2009a) captures snapshots of the long-range interactions of the genomic loci at the whole-genome level. Data from this technology consists of sequencing of millions of genomic locus pairs that are in physical contact and is summarized by a symmetric Hi-C contact matrix, entries of which represent a measure of physical contact between the locus pairs. Recent advancements in single cell sequencing technologies of Hi-C (scHi-C) enabled profiling interactions between distant genomic loci in individual cells (Stevens et al., 2017; Ramani et al., 2017; Tan et al., 2021; Ulianov et al., 2021) and even simultaneously with their DNA methylation status (snm3C-seq (Lee et al., 2019; Liu et al., 2021), scMethyl-HiC (Li et al., 2019a)). These new approaches have the potential to elucidate the interplay between the epigenetic mechanisms and 3D genome structure in a wide variety of biological contexts. Statistical and computational approaches for specific scHi-C data inference tasks are appearing rapidly (e.g., scHiCluster (Zhou et al., 2019), scHiC Topics (Kim et al., 2020), Higashi (Zhang et al., 2022a), BandNorm and 3DVI (Zheng et al., 2022), and Fast-Higashi (Zhang et al., 2022b), SnapHiC (Yu et al., 2021), scHiCTools (Li et al., 2021), DeTOKI (Li et al., 2021)). However, computational tools for integrating scHi-C with other data modalities such as transcriptomics, epigenomics, and epigenetics are lagging behind. Notably, the only method that can integrate scHi-C with scRNA-seq is scGAD (Shen et al., 2022). However, scGAD’s common feature-based integration approach does not capitalize on the simultaneous profiling of 3D conformation and DNA methylation status of cells as enabled by sn-m3C-seq (Lee et al., 2019; Liu et al., 2021) and scMethyl-HiC (Li et al., 2019a). In contrast, Higashi (Zhang et al., 2022a) facilitates joint analysis of scHi-C and DNA methylation data; however, the inference implemented is limited to cell type clustering and lacks downstream analysis, and its practical utility is hindered by its computational requirements, which led to development of Fast-Higashi (Zhang et al., 2022b). Fast-Higashi improved scalability of Higashi significantly; however, its current framework and implementation has not yet been leveraged to handle multiple modalities jointly.

In addition to the lack of integrative modeling approaches for scHi-C and DNA methylation, another key shortcoming of exiting scHi-C analysis methods, including scHiCluster, scHiC Topics, Higashi, 3DVI, and Fast-Higashi, is a lack of alignment between the parameters estimated by these methods and the key parameters of interest in the scHi-C analysis. While these methods are able to learn latent representations of individual cells for downstream cell type identification and clustering, inferring chromosome organization characteristics such as topologically associating domains (TADs) (Pombo and Dillon, 2015), A/B compartments (Lieberman-Aiden et al., 2009b) from their output requires, often complex, additional downstream processing of the estimated model parameters or denoised data after aggregating contact maps of the cells within each inferred cell type. From a strictly statistical perspective, the model parameters are estimated in isolation and without consideration of the intended post-processing procedures, and as a result, this might lead to unreliable and sub-optimal inference.

Here, we develop a new statistical model named Muscle for **Mu**ltiple **s**ingle **c**el**l** t**e**nsors to better align the estimated model parameters with the key inference parameters of scHi-C analysis and to enable integration of scHi-C with other data modalities. Muscle’s multiple tensor framework encodes parameters such as cell-specific loadings shared by all data modalities, loci loadings specific to data modalities, scHi-C eigen contact maps with one-toone alignments to the cell types, A/B compartment structures, loci groupings or TADs, cell type specific methylation profiles, all of which are critical for 3D genome and methylation analysis (**Fig**. 1). A key advantage of Muscle is that it can be deployed with only scHi-C data as well as with multiple single cell data modalities. Application of Muscle to multiple scHi-C datasets with ground truth (Lee et al., 2019; Kim et al., 2020; Ramani et al., 2017; Li et al., 2019b) demonstrates that Muscle performs as well or even better than existing methods for cell clustering and, more critically, can infer chromatin conformation structures in a cell type specific manner. Simulation studies comparing the joint analysis of Muscle to a baseline strategy of flatting out the tensor as a matrix and leveraging matrix decomposition reveal consistently better performance by Muscle and supports the robustness of Muscle to a wide range of signal-to-noise levels. Muscle in the joint analysis mode for the sn-m3C-seq data (Lee et al., 2019; Liu et al., 2021) or scMethyl-HiC data (Li et al., 2019a) successfully identifies cell type specific associations between DNA methylation profiles and 3D genome structure including TAD boundaries and compartment territories. Collectively, Muscle represents a significant modeling advancement in the joint analysis of scHi-C data with other data modalities.

**Figure 1:**
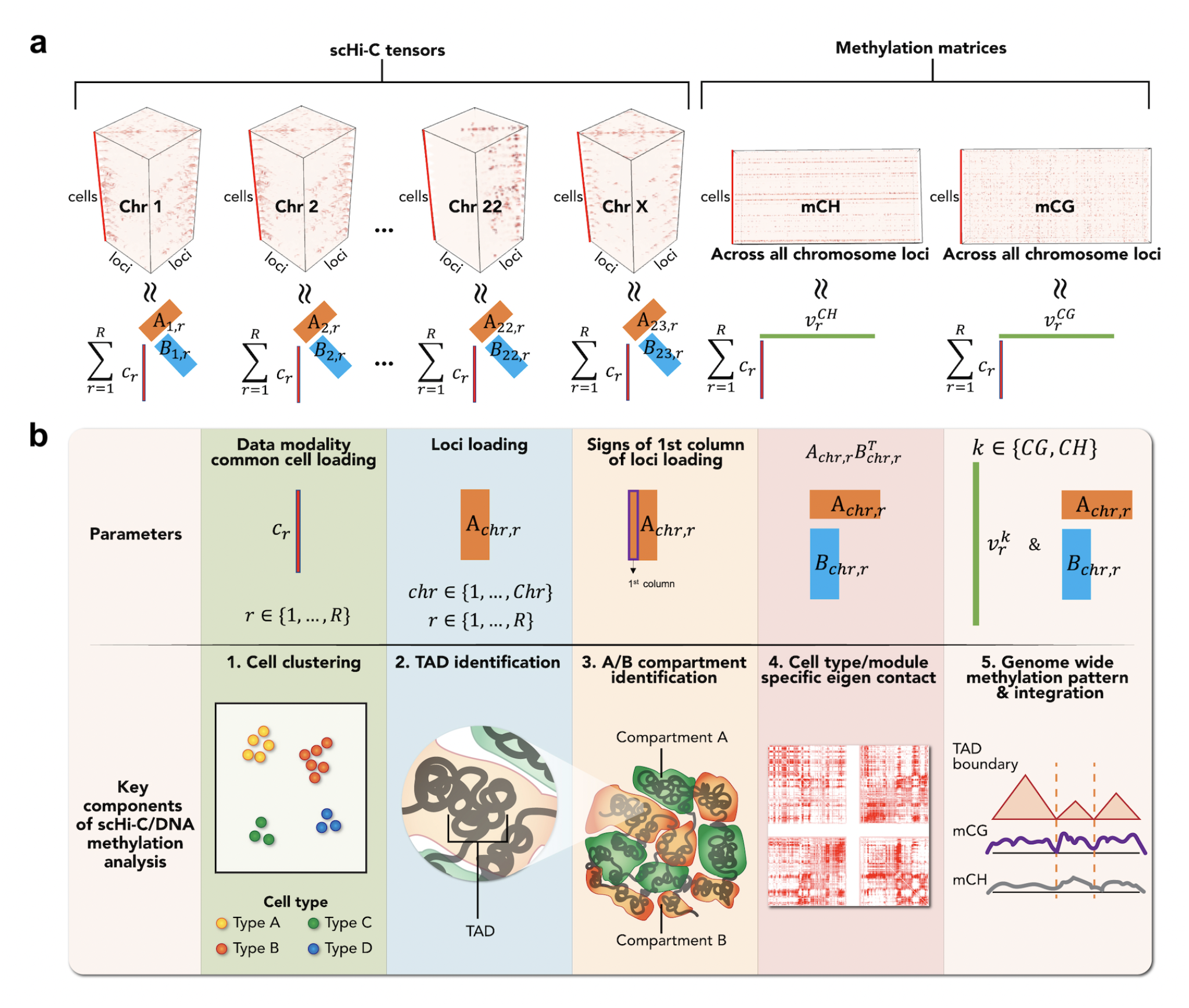
Overview of Muscle multiple single cell tensor model. **a**. Each chromosome-specific scHi-C tensor with size *l*_*chr*_ × *l*_*chr*_ × *C* is a summation of R “rank-1” modules 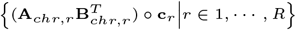, where *l*_*chr*_ and *C* denote the # of loci for chromosome *chr* and the # of cells, respectively. Each module contains three factor loadings. The data modality common cell loading **c**_*r*_ ≥ 0 encodes which cell type the module corresponds to and provides a “label/name tag” for the module. Each of the chromosome-specific loci loadings **A**_*chr,r*_, **B**_*chr,r*_ encompass structural chromatin characteristics of a specific cell type, and the eigen contact 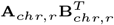 is the resulting interaction pattern (i.e., eigen contact map) of the cell type. Both mCG, mCH methylation matrices with size (∑_*chr*_ *l*_*chr*_) × *C* are also summation of “rank-1” modules 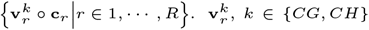, encodes the methylation profile along the genome for the cell type inferred from the cell loading **c**_*r*_. **b**. Muscle parameters align with the downstream analysis of interest. 1) The cell loading vectors {**c**_*r*_|*r* ∈ 1, …, *R*} enable cell clustering and identification of modules corresponding to each cell type. 2) Low dimensional projection of loci loading **A**_*chr,r*_ reveals loci clustering structure and TADs. 3) The first column vector of loci loading **A**_*chr,r*_ encodes A/B compartments which are large-scale genome territories. 4) Direct visualization of eigen contact map 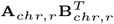 reveals contact pattern of the cell type that the *r*th module corresponds to. 5) The methylation loci loading vector 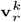 aligns with the eigen contact map 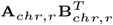 or scHi-C loci loading **A**_*chr,r*_ to yield associations between DNA methylation and 3D genome structure of the cell type identified by cell loading vector **c**_*r*_.

## 2 Muscle model

### 2.1 Muscle model representation

Muscle is a multiple tensor framework for single-cell multi-modal omics data. Here, we focus on scHi-C and DNA methylation and illustrate how Muscle parameters provide direct intuitive integrative inference of these single cell data modalities. **Fig**. 1a provides an overview of Muscle, which starts out with a tensor view of multi-modal scHi-C data and single cell DNA methylation data (top row). For scHi-C data modality, each set of *cis*-interaction (i.e., only intra-chromosomal interactions) contact matrices of a single chromosome is viewed as an order three tensor, with dimensions # of loci on the chromosome (denoted with *l*_*chr*_), # of loci on the chromosome, and # of cells (denoted with *C*). Here, each slice along the cell mode (or dimension) corresponds to a chromosome-specific scHi-C contact map for a single cell. For the human genome, this results in 23 order three tensors with the common cell mode but differing numbers of loci. For the single cell DNA methylation data, we form a mCG (mCH) matrix (i.e., order two tensor) with dimensions # of CpGs (non-CpGs) and # of cells, containing the CpG (non-CpG) site methylation level. The cell mode dimension is shared between the scHi-C and methylation tensors because of the multi-modality (i.e., scHi-C and methylation read outs are taken simultaneously from a single cell) of the data.

After forming the entire order two and three tensors, Muscle parameterizes each of their mean tensors (bottom row of **Fig**. 1a) following a semi-non negative Block Term Decomposition (BTD) form. Specifically, each of the scHi-C tensors is modelled as a summation of *R* “rank-1” modules, 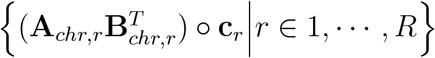. Note that we abuse the term “rank-1” to denote a rank-(*K*_*chr*_, *K*_*chr*_, 1) tensor for simplicity in this paper, where *K*_*chr*_ is defined as the block rank in BTD (De Lathauwer, 2008). Each rank-1 module captures a latent contact pattern of the data. The two chromosome-specific loci loadings **A**_*chr,r*_, 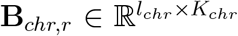 harbor physical interaction information of the module and a nonnegative data modality common cell loading vector (i.e., loadings shared by all data modalities) 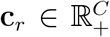 captures cell type information of the module. The methylation matrices are in the form of a semi-non negative matrix factorization, which is similarly a summation of *R* rank-1 modules 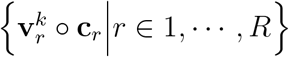, where *k* ∈ {*CG, CH*}. Likewise, the data modality common cell loadings **c**_*r*_, shared with scHi-C, encodes modules specific to cell types, and the methylation loci loadings 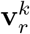 identify cell type specific methylation patterns.

Muscle formulation has two unique components that allow it to leverage multiple data modalities and enable direct inference for key parameters of interest. First, each cell loading vector **c**_*r*_ that is common to all data modalities (e.g., scHi-C and DNA methylation) learns the cell type information jointly across all chromosomes and data modalities, and, hence, is critical for the integrative analysis. Second, the non-negativity constraint on each cell loading vector **c**_*r*_ facilitates interpretation of each rank-1 module. For instance, if the cell loading vector of *r*th module has large values for a subset of the cells, i.e., from the same cell type, the matrix 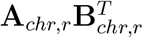 encodes the contact pattern for these groups of cells. In addition, the module’s loci loadings 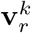 convey the cell type specific characteristics of genomic loci including A/B compartment structure and methylation profiles. We remark that in a PARAFAC2 (Kiers et al., 1999) decomposition-based model, which was utilized by the pioneering Fast-Higashi (Zhang et al., 2022b), similar interpretation is hindered by the sign indeterminacy issues of singular vectors, which are the “cell embeddings” of the Fast-Higashi. Specifically, aligning of modules with cell types can not be achieved if a cell embedding vector of the module has both large negative and positive values for different cell types. We discuss practical implications of this limitation in more detail in Supplementary Material Section section 4. In contrast, Muscle’s formulation achieves unification between model parameters and the key parameters needed for downstream inference. As a consequence, Muscle enables intuitive and direct interpretation of the model results as depicted in **Fig**. 1b. Each of Muscle’s model parameters or a combination thereof aligns with key inference parameters of 3D chromatin organization along with the

### 2.2 Statistical framework of Muscle

In this section, we introduce the statistical framework of Muscle and a brief overview of parameter estimation of Muscle in the next section. This exposition uses the following key definitions and notations. A set of sequential numbers, e.g., {1, …, *K*}, are abbreviated as [*K*]. For a third order tensor 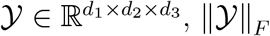 denotes the Frobenius norm. Finally, denotes outer product.

For a single cell *c* ∈ [*C*], we have *Chr* number of chromosomes and a symmetric Hi-C contact matrix of size *l*_*chr*_ × *l*_*chr*_ for each *chr* ∈ [*Chr*], where each (*i, j*)th entry of a contact map quantifies the observed physical interaction (e.g., contact) level between genomic loci *i* and *j*. For chromosome *chr*, the contact matrices stacked along cells have the same size *l*_*chr*_ × *l*_*chr*_. Hence, the data can be viewed as a (*l*_*chr*_, *l*_*chr*_, *C*)-dimensional tensor for each chromosome. We denote each pre-processed (e.g., natural log transformed for this specific setting) chromosome specific scHi-C tensor as 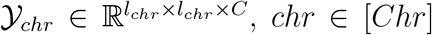. Further details about data pre-processing are discussed in Supplementary Material Section section 3.

The scHi-C tensors 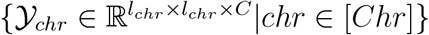 and methylation matrices **Y**^*CG*^, 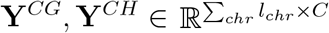, binned at 1Mb resolution (e.g., locus size), are modeled as

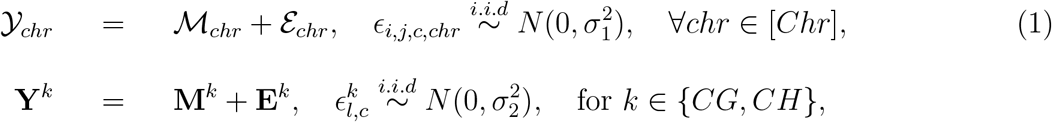

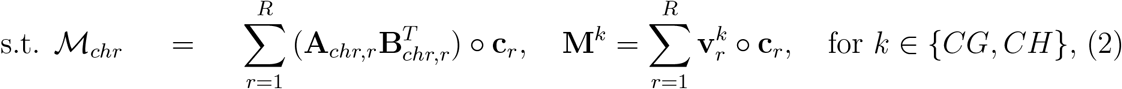

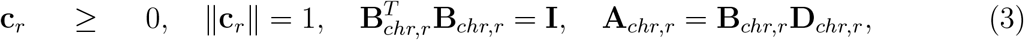

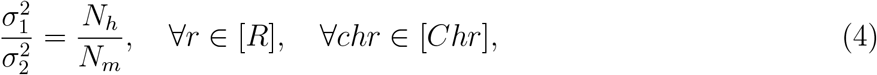

where all the chromosome specific signal and noise tensors 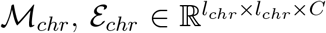, and mCG, mCH methylation signal and error matrices **M**^*CG*^, **M**^*CH*^, **E**^*CG*^, 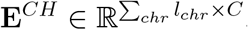. Also note that 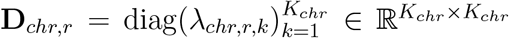 and *λ*_*chr,r,k*_ *>* 0 so that 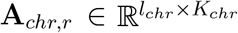 and 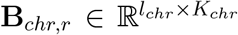 are equivalent up to multiplication of diagonal matrix absorbing the magnitude of the module. Here, the total size of scHi-C tensors is defined as 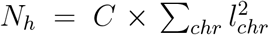, and, similarly, the size of a methylation matrix is defined as 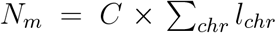. These size terms are leveraged to model the proportion of the variances of the two sources of data (Eqn. (4)). The signal tensor ℳ_*chr*_ is in the form of block term decomposition (De Lathauwer, 2008) and, the mean methylation matrices **M**^*CG*^, **M**^*CH*^ have the form of a semi-nonnegative matrix factorization. A key component of this integrative framework is that the non-negative cell loading vectors 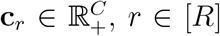, are shared between the methylation model and scHi-C tensor model, enabling the cell loadings to be learnt by leveraging both data modalities. We next provide an overview of parameter estimation for Muscle and refer to Supplementary Material Section section 1 for details.

### 2.3 Muscle model estimation

We introduce the estimation problem and a brief overview of the algorithm for Muscle. Given the scHi-C tensors 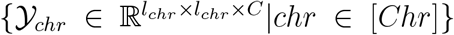 and methylation matrices **Y**^*CG*^, 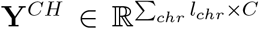, Muscle solves the following Maximum Likelihood Estimation equivalent problem

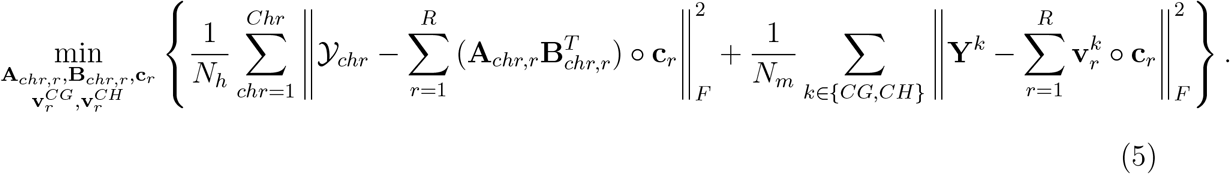

In order to solve this non-convex problem, we derive an Alternating Least Squares (ALS) algorithm (**Algorithm** 1 as described in the Supplementary Material Section section 1). The ALS algoritm iteratively obtains loci loading parameters **A**_*chr,r*_, **B**_*chr,r*_, and 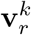 given the cell loadings **c**_*r*_ that are shared by both data modalities across all the chromosomes, and updates **c**_*r*_ by pooling all the loci loading information across the chromosomes. The estimation step of the cell loadings **c**_*r*_ is the key integrative part of Muscle. We derive the optimality properties of the ALS updates in Supplementary Material Section section 2.

## 3 Benchmarking with datasets with ground truth

### 3.1 Muscle is widely applicable for cell type identification with even single modality scHi-C data

We start out by exploring Muscle’s applicability with single modality scHi-C data by evaluating its cell clustering performance with multiple 3D genome datasets (Lee et al., 2019; Kim et al., 2020; Ramani et al., 2017; Li et al., 2019b). Details on parameter settings are provided in the Supplementary Material Section section 3. The ranks (*R*) of the low dimensional cell embeddings for Muscle and Fast-Higashi were set to be the same with the exception of the Lee et al. (2019) dataset which required a larger rank (*R* = 250) for Fast-Higashi to reveal cell types. Muscle enables cell clustering through the estimated cell loadings 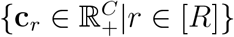. **Fig**. 2a,b display the scatter plots of the first two UMAP coordinates of cell embeddings from Muscle and other scHi-C analysis methods (Zheng et al., 2022; Li et al., 2021; Zhang et al., 2022a; Zhou et al., 2019; Kim et al., 2020) for the Li et al. (2019b) and Kim et al. (2020) datasets, which have the least and the most numbers of cells, respectively. Similar displays for rest of the datasets are available in Supplementary Material (**Fig**. S2). **Fig**. 2a highlights that Muscle and Higashi (Zhang et al., 2022a) are the only models that can separate Serum 1 cells from the others. Similarly, in **Fig**. 2b, the separation of the four major cell types (GM12878, H1Esc, HFF, HAP1) is more evident for Muscle, scHiC Topics (Kim et al., 2020), and 3DVI (Zheng et al., 2022) compared to the others. Next, we systematically evaluated the “cell type identification by clustering” performances of the methods. We used both the learned embeddings of the methods (e.g., cell loadings **c**_*r*_, *r* ∈ [*R*] for Muscle) and their low dimensional projections with UMAP and tSNE for cell clustering with *k*-means and employed the Adjusted Rand Index (ARI) and Average Silhouette Score metrics for evaluation based on ground truth cell labels (**Fig**. 2c). **Fig**. 2d summarizes the median ranking of the clustering performances for each of the combinations in **Fig**. 2c, and yields that Muscle shows the best ranking for cell clustering solely based on scHi-C data. This establishes Muscle’s applicability with scHi-C data even in the single data modality setting. In addition to these large-scale benchmarking experiments, we further explored the practical implications of the differences in the formulations of Muscle and PARAFAC2-based Fast-Higashi on the Kim et al. (2020) scHi-C dataset in more depth in Supplementary Section section 4.

**Figure 2:**
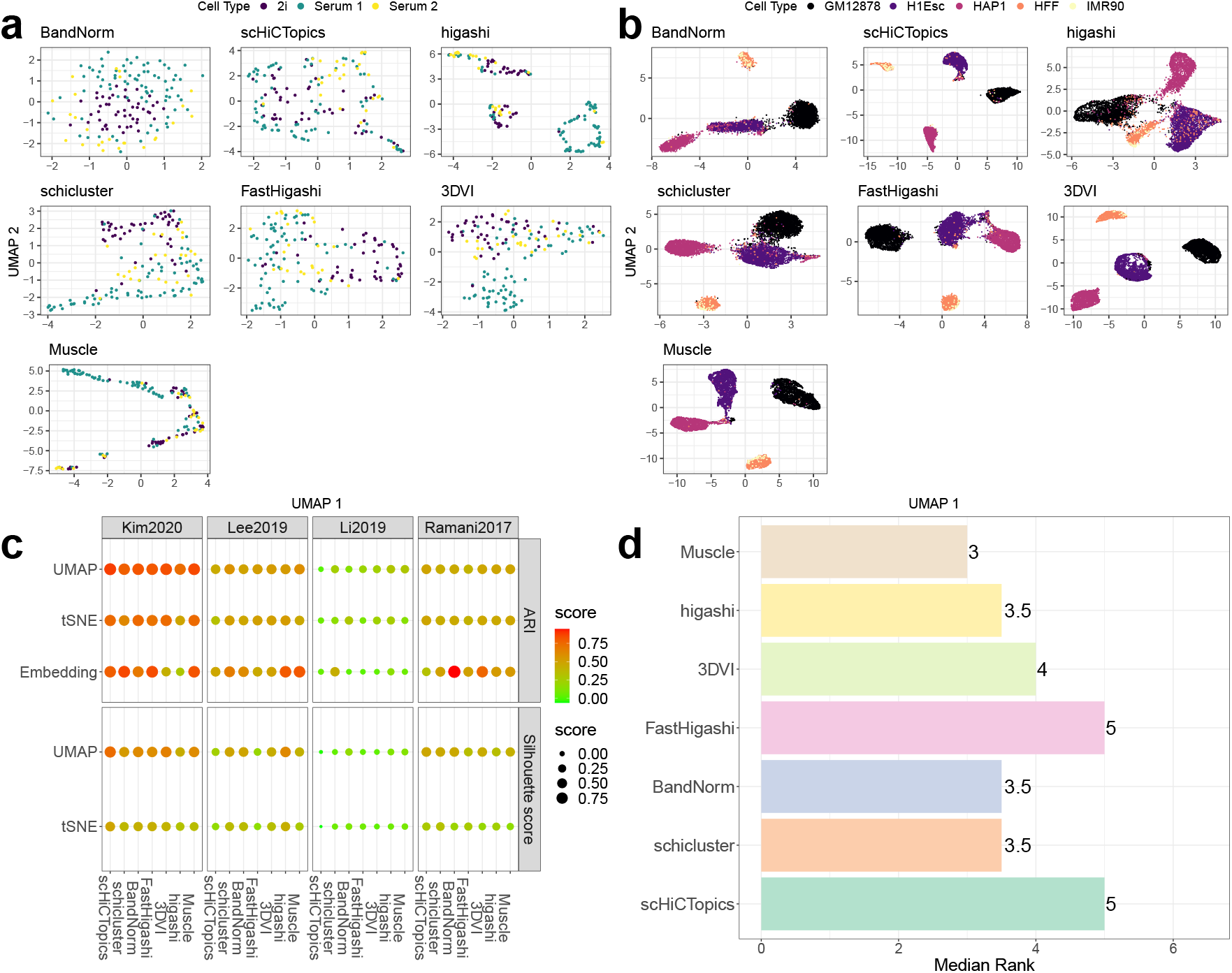
Computational evaluation and benchmarking of Muscle cell clustering with scHi-C data. **a, b**. UMAP coordinates of the cells from Li et al. (2019b) and Kim et al. (2020) scHi-C datasets. Muscle UMAP coordinates are obtained from estimated 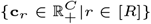. Cells are colored based on known cell type labels. **c**. Evaluation of the cell clustering by different methods. Larger and redder circles correspond to larger scores. **d**. Median ranking of the methods across the multiple evaluation settings in panel **c**.

### 3.2 Integrative framework of Muscle improves cell type clustering in the multi-modal setting

After establishing Muscle’s on par performance with existing methods in the single modality setting, we turned our attention to the integrative framework. We utilized the Lee et al. (2019) and Liu et al. (2021) sn-m3C-seq datasets, which simultaneously profiled 3D genome and DNA methylation in 14 human brain prefrontal cortex cell types and 10 mouse hippocampal cell types, respectively. In the integrative analysis, the cell loadings 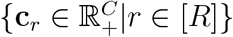 are learnt utilizing both data modalities. **Fig**. 3a,c provide a direct comparison of matrix factorization via singular value decomposition (SVD) using only DNA methylation components (only mCG or only mCH; top middle, top right panel of **Fig**. 3a,c) and Muscle using only the scHi-C (top left panel of **Fig**. 3a,c) with the integrative Muscle (bottom right panel of **Fig**. 3a,c). Visual inspection of **Fig**. 3a,c reveal how Muscle leverages different data modalities. Specifically, for Lee et al. (2019) data, the integrative model (bottom right panel of **Fig**. 3a) provides complete separation of the inhibitory neuoronal cell types (Ndnf, Pvalb, Sst, Vip cells), while the results for scHi-C only and mCG only modalities lack such a separation (top left, top middle). The separation of excitatory cells (L23, L4, L5, L6 cells) for multi modal Muscle is as evident as in the result from mCG only case (top middle), and markedly more apparent than that of the scHi-C only case (top left). While the OPC and ODC cells are not separated in mCG and mCH only modalities (top middle, top right), these cells can be separated in the integrative analysis (bottom right). We also compared the Muscle cell clustering results in the multi-modal setting (**Fig**. 3a bottom right) with a baseline approach (**Fig**. 3a bottom left) that performed principal component analysis on the matricized “*cell* × *locus pair* “ scHi-C data concatenated with mCG, mCH matrices. This approach used the same rank as the Muscle rank *R*, and the resulting *R* principal components were utilized to generate the UMAP embeddings. The bottom left panel of **Fig**. 3a shows that the cell type separation from this baseline approach is inferior to that of Muscle depicted in **Fig**. 3a. Moreover, the baseline approach is susceptible to the batch effects in the excitatory and inhibitory neuronal cell types, whereas the Muscle result is relatively free of this effect (**Fig**. S3a). The overall performances of these settings, which reveal the marked improvement by the Muscle multi modal setting, are summarized in **Fig**. 3b.

**Figure 3:**
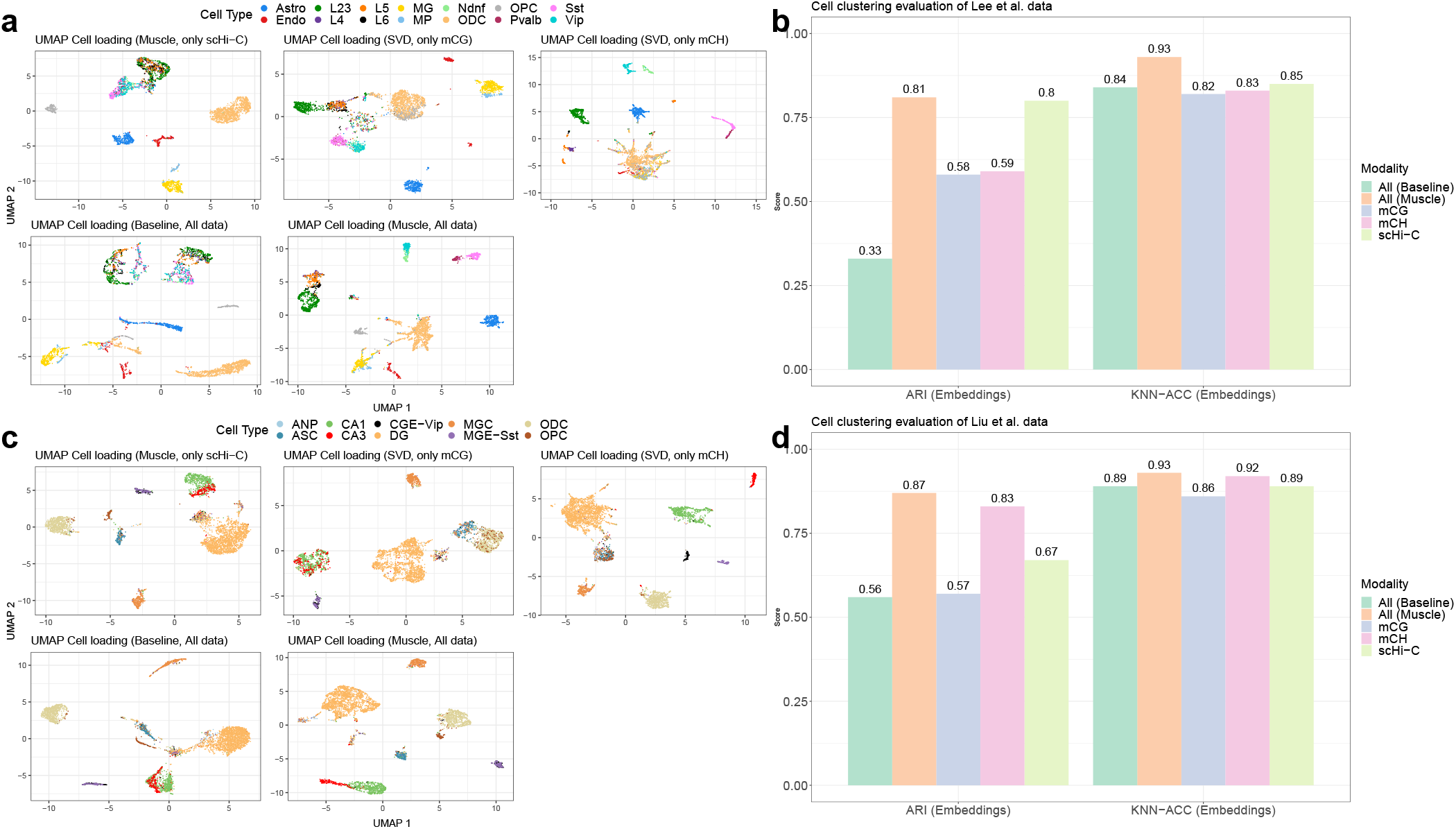
Computational evaluation and benchmarking of Muscle cell clustering with the multi-modal set up. **a**. (Top-left) UMAP coordinates of the cells from Lee et al. (2019) sn-m3C-seq data based only on scHi-C modality. Muscle UMAP coordinates are obtained from estimated cell loading vectors 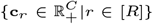. Cells are colored based on known cell type labels. (Top-middle) UMAP coordinates of the cells from Lee et al. (2019) sn-m3C-seq data based only on the mCG methylation modality. The UMAP coordinates are obtained from estimated cell loading vectors 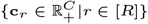 of SVD. (Top-right) UMAP coordinates of the cells based only on mCH methylation modality. (Bottom-left) UMAP coordinates of the cells based on both scHi-C and DNA methylation modalities, obtained from baseline matricization method. The UMAP coordinates are obtained from principal components of the concatenated (matricized) scHi-C and DNA methylation datasets. (Bottom-right) UMAP coordinates of the cells based on both scHi-C and DNA methylation modalities, obtained from Muscle. **b**. Left : The ARI scores from *k*-means clustering of the cells with the learned embeddings under single (scHi-C only, mCG only, mCH only) and multi-modal settings (Muscle, baseline). Right : K Nearest Neighborhood (KNN) classification accuracy with cell loadings as features and ground truth cell type labels as classes. KNN results are averaged over 20 sets of training-test data splits where test data harbored 10% of the randomly selected cells. The number of neighbours was set as *K* = 20. **c**. (Top-left) UMAP coordinates of the cells from Liu et al. (2021) sn-m3C-seq data based only on scHi-C modality. Muscle UMAP coordinates are obtained from estimated cell loading vectors 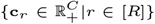. Cells are colored based on known cell type labels. (Top-middle) UMAP coordinates of the cells from Liu et al. (2021) sn-m3C-seq data only on the mCG methylation modality. The UMAP coordinates are obtained from estimated cell loading vectors 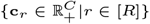 of SVD. (Top-right) UMAP coordinates of the cells based only on mCH methylation modality. (Bottom-left) UMAP coordinates of the cells based on both scHi-C and DNA methylation modalities, obtained from baseline matricization method. The UMAP coordinates are obtained from principal components of the concatenated (matricized) scHi-C and DNA methylation dataset. (Bottom-right) UMAP coordinates of the cells based on both scHi-C and DNA methylation modalities, obtained from Muscle. **d**. Left : The ARI scores from *k*-means clustering of the cells with the learned embeddings under single (scHi-C only, mCG only, mCH only) and multi-modal settings (Muscle, baseline). Right : K Nearest Neighborhood (KNN) classification accuracy with cell loadings as features and ground truth cell type labels as classes. KNN results are averaged over 20 sets of training-test data splits where test data harbored 10% of the randomly selected cells. The number of neighbours was set as *K* = 20.

Analysis of a more recent sn-m3C-seq dataset from mouse hippocampus (Liu et al., 2021) provided insights similar to those of the above analysis. Specifically, results from the analysis of this dataset revealed that the integrative Muscle enabled complete separation of the cell types CA1 and CA3 (**Fig**. 3c bottom right). These cell types appeared to be less separated in the scHi-C only and mCG only analysis (**Fig**. 3c top left and middle). In this case, Muscle leveraged the cell type separation information of CA1 and CA3 cells from the mCH modality (**Fig**. 3c top right). Similarly, while delineation of the cell type ASC from only the mCH modality exhibited ambiguity (**Fig**. 3c top right), the integrative Muscle model achieved good separation of this cell type from the others (**Fig**. 3c bottom right) by leveraging the cell type separation information from the scHi-C modality (**Fig**. 3c top left). The overall performances of these settings, which reveal the marked improvement by the Muscle multi modal setting over integrative baseline method and single modalities, are

### 3.3 Muscle yields cell type specific modules that delineate cell type specific contact maps

Next, we explored the inference readily available from Muscle for downstream scHi-C analysis with the Kim et al. (2020) data, containing five human cell lines (GM12878, H1Esc, HAP1, HFF, and IMR90). Muscle’s rank-1 modules 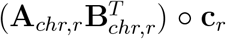 capture parsimonious representations of the cell types. The magnitudes of the Muscle cell loading vectors, e.g., **c**_*r*_ ≥ 0, *r* = 1, …, *R*, delineate cells corresponding to the same cell type/state, therefore linking modules to specific cell groups. Consequently, the corresponding loci loadings, 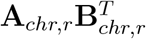, for module *r* describe the denoised contact map of the corresponding cell type. For example, Muscle cell loading vector *c*_7_ has exclusively large values for the GM12878 cells (**Fig**. 4b). Hence, the eigen contact map 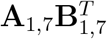 depicted in **Fig**. 4e corresponds to the mean contact pattern of chr 1 in GM12878 cells. Similarly, 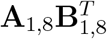 in **Fig**. 4f displays the HFF specific contact pattern because the cell loading vector *c*_8_ of this module is specific to cell type HFF, i.e., with large positive entries for HFF cells (**Fig**. 4c). In addition, notice that *c*_1_ has constant values throughout cells (**Fig**. 4a). Hence, this module can be interpreted as a grand mean pattern of the entire cell types, and the first module’s eigen contact map 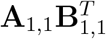 depicted in **Fig**. 4d corresponds to the grand average contact map of all the cell types.

**Figure 4:**
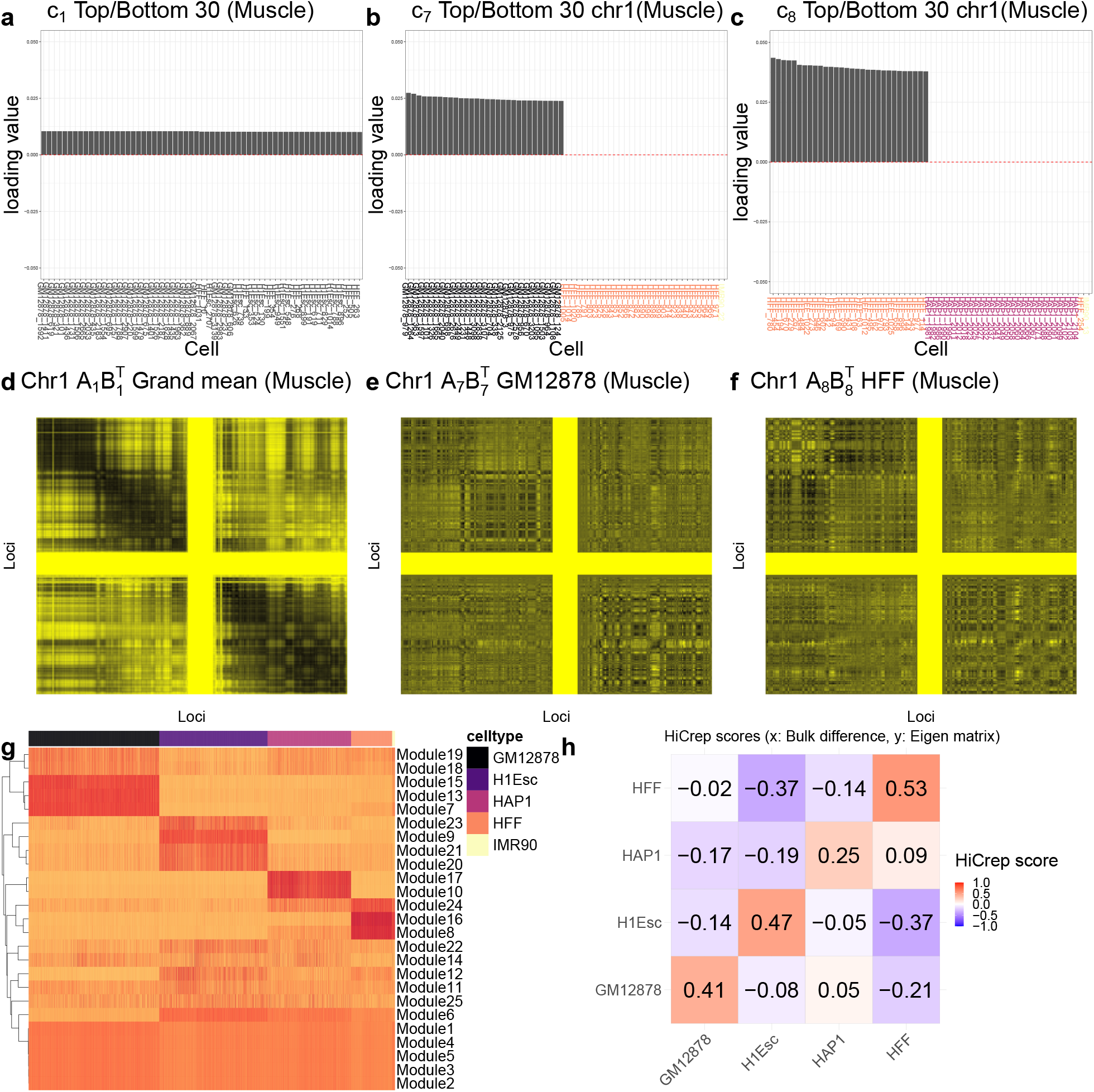
Cell type specific eigen-contact captures by Muscle. **a, b, c**. Muscle cell loading vectors *c*_1_, *c*_7_, *c*_8_ depicted as barplots, respectively. The cell loading vectors are constrained to be non-negative and yield the representative cell type of each module. X-axis labels are colored based on the true cell type labels of the cells in panel g. Top and bottom 30 cell loadings are displayed for brevity. **d, e, f**. Visualization of Muscle eigen contact maps (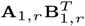terms) for *r* = 1, 7, 8, which capture grand average, GM12878 and HFF contact patterns. In panels d,e,f, darker entries indicate higher interactions between the loci. **g**. Heatmap of the entire set of Muscle cell loading vectors 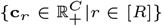. Each row displays the estimated cell loadings **c**_*r*_ of the module and each column corresponds to a cell. **h**. HiCrep score comparison of Muscle eigen contact maps 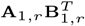 of modules *r* = 7 (GM12878), *r* = 8 (HFF), *r* = 9 (H1Esc), and *r* = 10 (HAP1) against the gold standard cell type specific bulk contact maps. The y-axis denotes inferred cell type specific eigen contact maps and the x-axis denotes the cell type specific bulk contact maps.

To validate that the eigen contact maps, 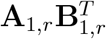, are indeed cell type specific, we calculated HiCRep similarity scores (Yang et al., 2017), a modified version of Spearman correlation to compare two Hi-C contact matrices, between cell type specific contact maps (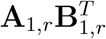, where different *r* values correspond to different cell types) and cell type specific contact maps generated from the gold standard cell type specific bulk data (Kim et al., 2020). The heatmap in **Fig**. 4g displays the entire cell loading vectors **c**_*r*_ and clearly demarcates modules specific to each cell type. While several cell types have multiple modules, we chose *r* ∈ {7, 8, 9, 10}, each of which had the largest size ∥**A**_1,*r*_∥_*F*_ (i.e., parameters with the largest effect sizes) among the modules corresponding to the cell types GM12878, H1Esc, HFF, and HAP1, respectively. **Fig**. 4h demonstrates that the similarity score is the highest when the eigen contact map 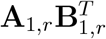 of a cell type is compared against its own gold standard bulk data (i.e., largest similarity scores along the diagonal). Collectively, these results further proffer the main advantages of Muscle’s tensor decomposition framework which targets the key parameters.

### 3.4 Muscle yields cell type specific TADs and A/B compartments

Topologically associating domains (TADs) constitute large genomic regions with larger numbers of interactions between loci within the region compared to interactions of loci with the loci outside the region. TADs are highly cell type specific since they recapitulate cell type specific regulation (Yu and Ren, 2017). Muscle parameters **A**_*chr,r*_ reveal TADs for module *r*, and the cell type of the module is delineated by the positive entries of **c**_*r*_. We note that the loci loading **A**_*chr,r*_ is used instead of **B**_*chr,r*_ for elucidating TADs. These two are identical up to module magnitude multiplication because of the symmetry of the cell contact matrices, and **A**_*chr,r*_ is more appropriate for inference since it absorbs the magnitude of the module *r* (see Eqn. (3)).

We next investigate the TADs for the Kim et al. (2020) scHi-C data analyzed in the previous section. **Fig**. 5a displays the UMAP plot of loci loadings **A**_1,8_ of module 8, which corresponds to cell type HFF (as depicted in **Fig**. 4b), for chr 1. Coloring the loci in this plot according to the TADs identified from the gold standard HFF bulk data (Supplementary **Fig**. S4) reveals that the loci loading **A**_1,8_ of Muscle organizes the loci within a chromosome in a way consistent with their TAD structures.

**Figure 5:**
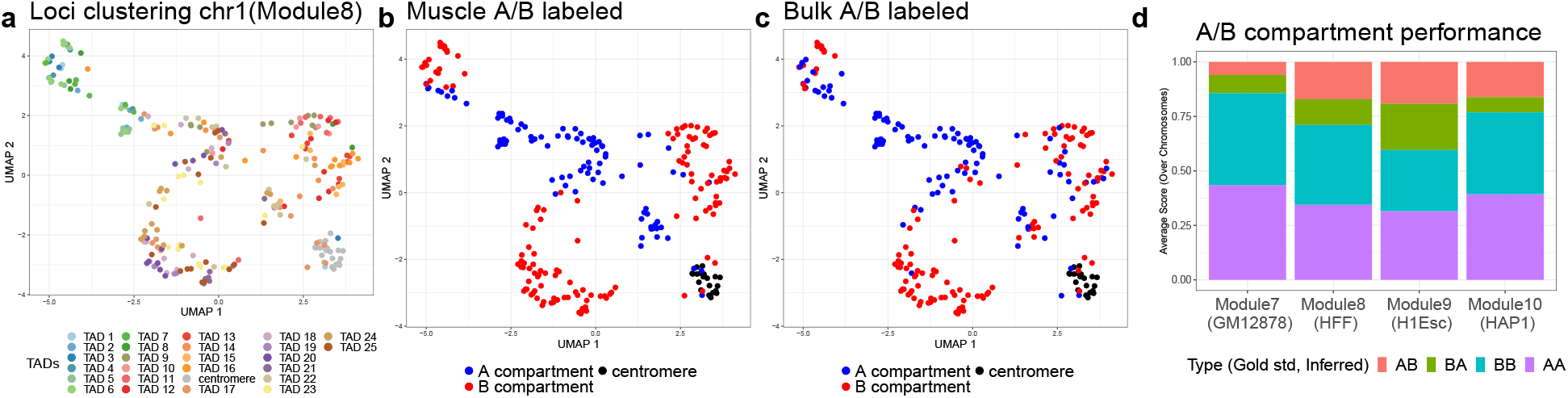
TAD and A/B compartment identification from estimated Muscle parameters. **a**. UMAP coordinates of Muscle estimated **A**_1,8_ of (Kim et al., 2020) dataset. Each data point corresponds to one locus of chr 1 and the colors are based on the gold standard TADs from the bulk HFF Hi-C contact map depicted in **Fig**. S4. **b**.UMAP coordinates of chr 1 loci from panel (a) colored with respect to the signs of the first column of **A**_1,8_, with “+” depicted in blue and corresponding to A compartment and “-” depicted in red and corresponding to B compartment. **c**. UMAP coordinates of chr 1 loci from panel (a) colored with respect to the gold standard A/B compartment results from HFF bulk Hi-C data. **d**. Evaluation of A/B compartment territory inference of Muscle based on the gold standard A/B compartments from cell type specific bulk Hi-C data. Each barplot represents a cell type (module) and displays the mean proportion of correctly inferred A (or B) compartments averaged across the chromosomes. The first element of a label (e.g., AB) is for true compartment and the other element is for inferred compartment. The regions corresponding to the centromere are excluded from the analysis.

A/B compartments, which constitute genome territories with high (A) or low (B) gene expression compared to other territories, is another class of genome compartmentalization that can be inferred from scHi-C data (Lieberman-Aiden et al., 2009b). In addition to identifying TADs, the first column vector of **A**_*chr,r*_ for each *chr* ∈ [*Chr*] and *r* ∈ [*R*] provides the A/B compartment structures in a cell type specific manner. This is because the loci loadings **A**_*chr,r*_, **B**_*chr,r*_ are obtained from an eigen decomposition of the projected scHi-C tensor 𝒴_*chr*_ onto the subspace spanned by **c**_*r*_ (**Algorithm** 1), and hence, the first column of **A**_*chr,r*_ is formed by the multiplication of the largest eigenvalue and the corresponding eigenvector. As a result, the first column of **A**_*chr,r*_ captures the major contact pattern of the module *r*, which would represent the largest scale genome territory, i.e., A/B compartments. **Fig**. 5b displays the exact same loci clustering of **A**_1,8_ as in **Fig**. 5a where the labels of loci are now obtained by the sign of the first column of **A**_1,8_. The interior loci in **Fig**. 5b are inferred to be in A compartments, while the loci in the exterior are in the B compartments. We validated this labeling by comparing it to the labeling from the gold standard HFF bulk Hi-C data’s A/B compartment structure (**Fig**. 5c). We further evaluated the A/B compartment inference for each cell type (as identified by modules *r* ∈ {7, 8, 9, 10}) by comparing inferred compartmentalizations (averaged over all the 23 chromosomes) with those from the cell type specific bulk data (**Fig**. 5d). This evaluation yielded that, on average, 72% and 75% of the true A and B compartment loci were correctly identified, respectively, by Muscle. In summary, the estimation targets of Muscle directly matches the parameters of interest in scHi-C data analysis and the estimated parameters readily reveal TAD and A/B compartment structures of the cell types without additional downstream analysis.

### 3.5 Muscle unveils cell type specific associations between chromatin conformation structures inferred from scHi-C data modality and DNA methylation modality

DNA methylation in both the CpG and non-CpG sites is generally negatively correlated with the gene expression levels in mammallian neuronal cells (Luo et al., 2018; Lister et al., 2013). Using Muscle integrative analysis results of Lee et al. (2019) sn-m3C-seq data, we explored whether a given module’s methylation loci loadings 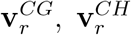 associated with the A/B compartment structure of the loci inferred from scHi-C modality loci loadings **A**_*chr,r*_[, 1]. Specifically, we investigated the association between genome compartmentalization and DNA methylation pattern in an excitatory neuronal cell type (L5) and in an inhibitory neuronal cell type (Vip). These two cell types are shown to be well-separated in the integrative Muscle analysis (bottom right panel of **Fig**. 3a). We observed that, as expected, the mCG and mCH methylation loci loadings 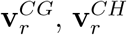 displayed show negative association with scHi-C loci loadings **A**_1,*r*_[, 1], where *r* = 15 and *r* = 21 denote cell types L5 (**Fig**. 6a) and Vip (**Fig**. 6d), respectively. Formal evaluation of association between fitted methylation (aggregated over a cell type) and A/B compartmentalization confirmed this observation (**Fig**. 6b, e) and revealed that the association is cell type specific. In Vip cells, loci in B compartment territory have significantly higher methylation levels on CpG sites than those of the loci in the A compartment (*p* = 9.7 × 10^−6^), while the association in L5 cells was not as significant as that of Vip cells (*p* = 0.07). Similarly, the differences of non-CpG methylation over the A and B compartments varied in a cell type specific manner as well. For Vip cells, non-CpG methylation over the A and B compartments did not exhibit statistically significant differences (*p* = 0.34, right panel of **Fig**. 6e). However, for L5 cells, non-CpG methylation differed between A and B compartments more than that of the CpG methylation (*p* = 0.031, **Fig**. 6b). These reinstate that association of loci methylation levels with genome territorial structures is cell and methylation site type (CpG or non-CpG) specific.

**Figure 6:**
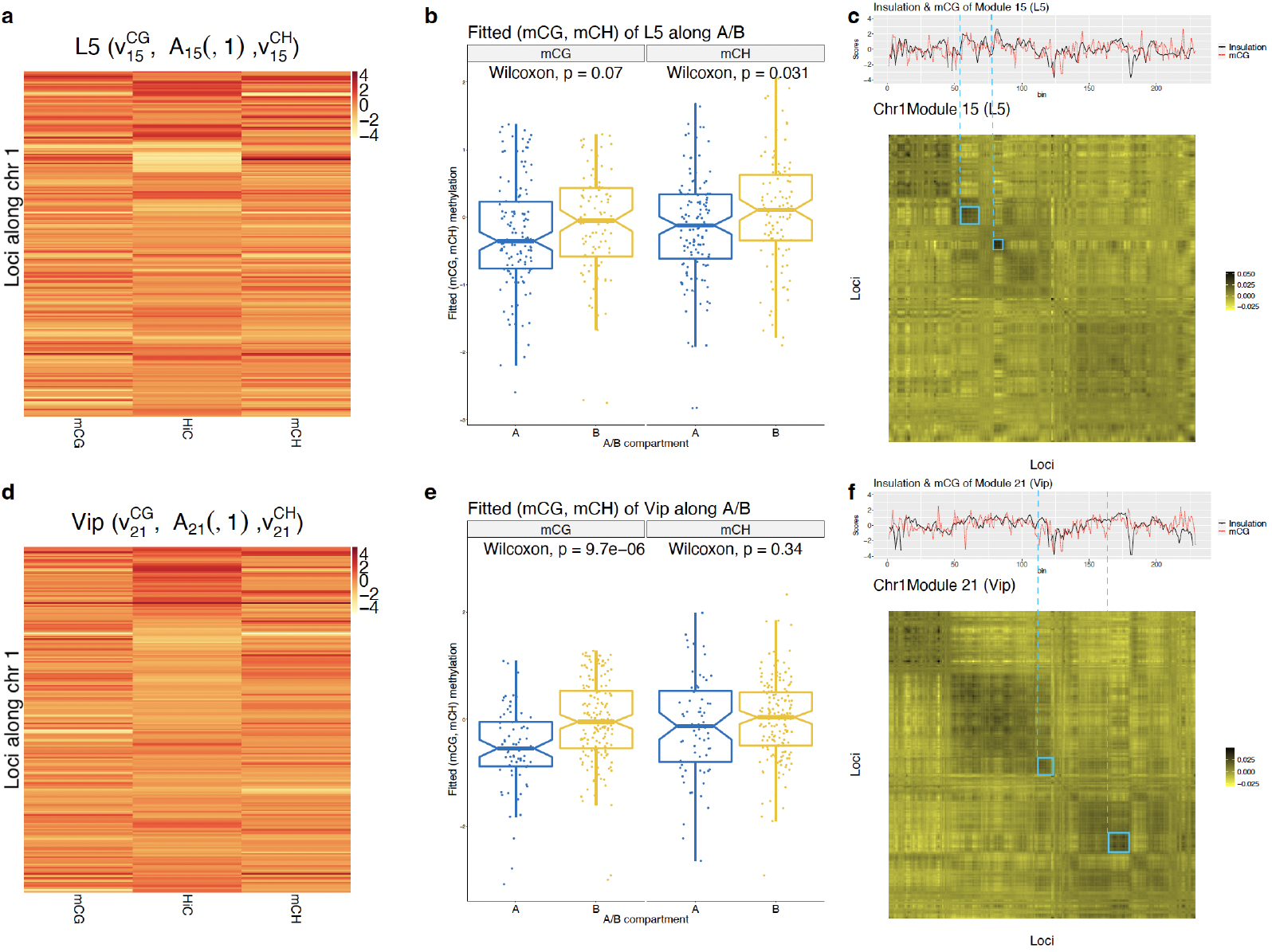
Association analysis of methylation patterns with broader 3D chromatin structures. **a, d**. Heatmap displays of the: (left) mCG methylation loci loading 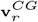 along chr 1 loci (*r* = 15 for panel (a) and *r* = 21 for panel (d)), (middle) the loci loading vector **A**_1,*r*_ [, 1] of chr 1 from the scHi-C part of the model, (right) mCH methylation loci loading 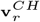 along chr 1 loci (*r* = 15 for panel (a) and *r* = 21 for panel (d)). Y-axis denotes the genomic loci of chr 1. **b, e**. Boxplots of fitted methylation levels averaged over cell type L5(b) and Vip(e) stratified with respect to CpG (mCG) and non-CpG (mCH) and within A/B compartments for L5 (panel b, identified from **A**_1,15_ [, 1]) and Vip (panel e, identified from **A**_1,21_ [, 1]) cells. Differences in methylation levels are evaluated with a Wilcoxon rank-sum test. **c, f**. Top row of the panel: the dotted red line displays the inferred methylation pattern along chr 1 loci, scaled as 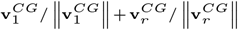. The solid black line represents insulation scores obtained from scaled eigen contact maps, 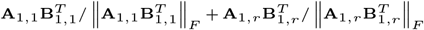. Bottom rows of the panels display scaled eigen contact maps. Note that cell type specific eigen contact maps are obtained by aggregating scaled eigen contact maps of modules common to all cell types (e.g., module *r* = 1 for this application) and cell type specific modules: 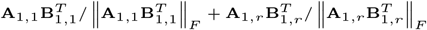 where *r* = 15 (cell type L5) in panel **c** and *r* = 21 (cell type Vip) in panel **f**.

Finally, we exploited the integrative analysis results from the point of CCCTC-binding Factor (CTCF) DNA binding protein. The activities of CTCF are inhibited by DNA methylation around the CTCF binding sites (Wang et al., 2012). In particular, DNA methylation plays a significant role in disruption of CTCF binding around key tumor suppressing genes in cancer (Rodriguez et al., 2010). CTCF is also a key player in folding of chromatin into domains. Specifically, TAD boundaries, where cohesin and CTCF form a DNA binding complex to hold the DNA loops together, are enriched for CTCF binding sites (Pombo and Dillon, 2015; Rao et al., 2014). As a result, we expect that the TAD boundary regions have more CTCF binding, and hence less DNA methylation that would hinder CTCF binding activity. **Fig**. 6c, f display methylation patterns from the methylation loci loading parameter 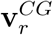 and the insulation scores (Gong et al., 2018), which quantify how *unlikely* a locus is to be a TAD boundary, from the eigen contact map 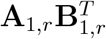, together with the display of the corresponding eigen contact maps. These comparisons reveal that insulation score patterns align well with the large-scale methylation patterns for both cell types. It further corroborates that genomic loci that are likely to be TAD boundaries (i.e., with low insulation scores marked with the blue dashed lines) have low methylation levels.

## 4 Simulation studies

Datasets with known cell types enabled us to illustrate the superior performance of the joint analysis with Muscle against both the single modality analysis and a baseline integrative approach. We further studied advantages of the tensor decomposition framework of Muscle over the matricization-based baseline method with simulation experiments. In these experiments, we ensured that the data generation process does not conform with Muscle’s model (given in Eqn. (1)-(4)) to quantify Muscle’s robustness against model misspecification.

### 4.1 Data generation

The scHi-C tensor 𝒴 ∈ ℝ^40 × 40 × 120^ and the DNA methylation matrix *Y* ∈ ℝ^40 × 120^, for 40 genomic loci and 120 cells across three cell types (with 40 cells from each cell type), were simulated from the following Negative binomial models:

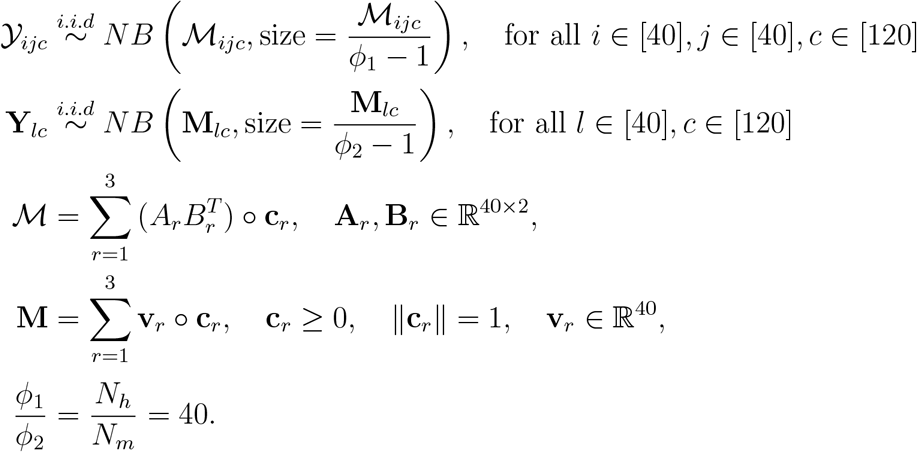

We generated three cell types by setting the entries 1 to 40 of *c*_1_, 41 to 80 of *c*_2_, and 81 to 120 of *c*_3_ to 3.1 and all the other entries of the cell loading vectors to 1 before size normalization. For each module *r, r* ∈ [3], the first column of loci loading matrix **A**_*r*_ ∈ ℝ^40×2^ is set to represent the A/B compartment structure (checker board-like pattern) of a contact map and the other column is set to represent a single TAD structure (square box-like pattern) along the diagonals of a contact map. The scHi-C loci loading matrix **B**_*r*_ ∈ ℝ^40×2^ is a column-wise normalization of **A**_*r*_ so that it becomes equivalent to **A**_*r*_ up to module size magnitude. This formulation, in turn, generates each eigen contact of the contact map 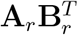 as in **Fig**. S6a-c. The methylation loadings **v**_*r*_ are randomly generated from a Poisson distribution with rate parameter *λ* = 0.23. The constructed methylation modules **v**_*r*_ ° **c**_*r*_ are displayed in **Fig**. S6j-l. The distributional assumptions on 𝒴 and *Y* result in

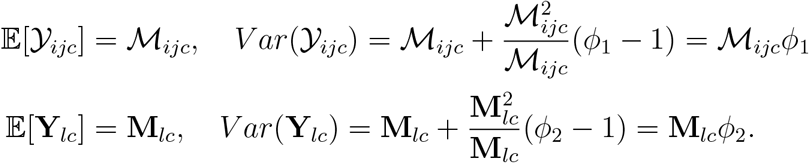

This data generation set up further ensures that ℳ_*ijc*_ ≈ **M**_*lc*_ for all *i, j, l, c* (the difference between the medians is *<* 0.2, and the difference between the means is *<* 0.2) as depicted in **Fig**. S7e. Consequently, the proportion of the variances between two data modalities approximately satisfies

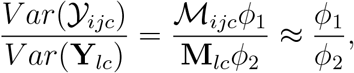

and allows us to vary the proportion of the variances of the two sources of data by modulating *ϕ*_1_, *ϕ*_2_. Under this data generation scheme, the resulting scHi-C and DNA methylation data exhibit general characteristics of the observed scHi-C and DNA methylation datasets (examples are provided in **Fig**. S7a-d).

### 4.2 Simulation results

We varied the noise level of the methylation data as *ϕ*_2_ ∈ {1.1, 1.2, …, 3} and the noise level of the scHi-C data *ϕ*_1_ is automatically determined based on the proportional variance construction as described in the previous section. For each of the *ϕ*_2_ values, we generated 30 simulation replicates and quantified the performances of Muscle and the matricization-based baseline method across these replicates. For each Muscle fit, the proportion of the variance was set to 40. The rank *R* was set *R* = 4 for both methods.

Comparison of the cell clustering performances of the methods revealed that Muscle results in significantly higher ARI scores than the baseline method, with a median difference of 0.05 across all the noise levels, *ϕ*_2_ (**Fig**. 7a,b, *p* = 0.043 based on Wilcoxon rank-sum test of the ARIs of the two methods).

**Figure 7:**
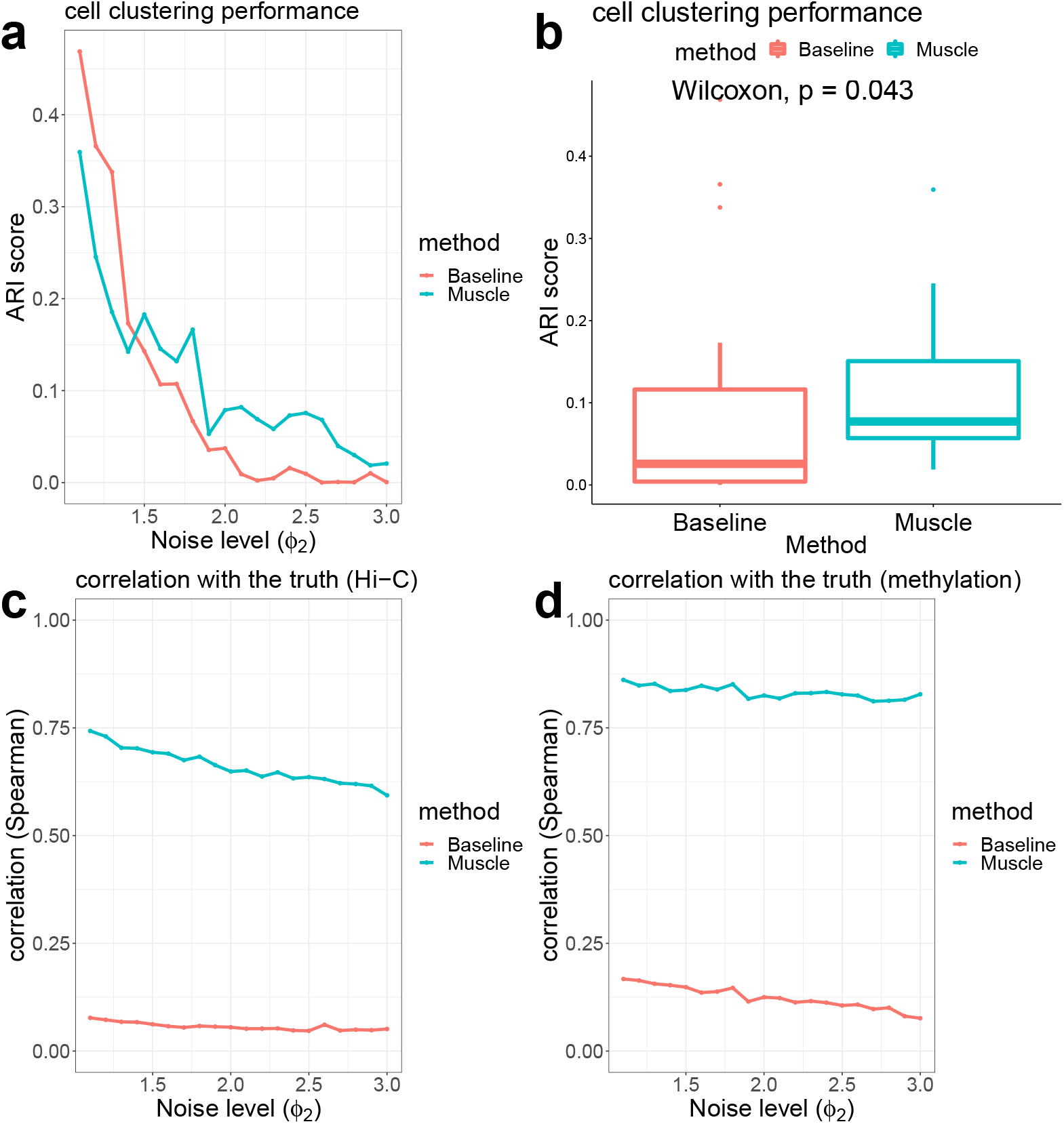
Evaluation of multi-modality analysis by Muscle and the matricization-based baseline method with simulation studies. **a**. ARI scores of the methods across all the noise levels *ϕ*_2_ averaged over 30 replicates. **b**. ARI scores of the baseline and Muscle results across all the noise levels, *ϕ*_2_. Results for fixed *ϕ*_2_ are averaged across the simulation replicates. Differences in ARI scores are evaluated with a Wilcoxon rank-sum test. **c**. Spearman correlations of the methods between the true scHi-C mean tensor ℳ and the estimated mean tensor 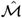 across noise levels *ϕ*_2_ averaged over 30 replicates. **d**. Spearman correlations of the methods between the true mean DNA methylation matrix *M* and the estimated mean methylation matrix 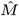 across noise levels *ϕ*_2_ averaged over 30 replicates.

We next investigated how well each method recovers the true means ℳ and **M**. Specifically, we evaluated the Spearman correlation between the true ℳ and estimated 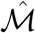 for the scHi-C modality and the Spearman correlation between the true **M** and estimated 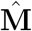 for the DNA methylation modality for both of the methods. **Fig**. 7c illustrates that the estimates from the baseline model have almost zero correlations with the true mean scHi-C contact matrices, whereas Muscle estimates have markedly higher correlation values (≈ 0.7) across all the noise levels *ϕ*_2_. This is also evident visually from **Fig**. S6. While the baseline method results in markedly noisy eigen contacts (**Fig**. S6g-i), the Muscle eigen contacts reasonably capture the cell type specific modules (**Fig**. S6d-f). Likewise, **Fig**. 7d also illustrates that the baseline model results in low correlations with the true mean DNA methylation matrix (≈ 0.15), while Muscle achieves markedly higher correlation values (≈ 0.8) across all the noise levels *ϕ*_2_. This can also be visualized in **Fig**. S6j-r, which highlights that the modules derived from the baseline method is not denoised enough compared to those from Muscle.

More detailed results on the cell clustering and the recovery of the mean scHi-C tensor ℳ and mean methylation matrix **M** are provided in **Fig**. S8. These specifically summarize the results based on the setting with the noise level *ϕ*_2_ = 1.1. The UMAP plots of the cell loadings depicted in **Fig**. S8a-b show that Muscle exhibits more apparent cell clustering than the baseline method (clustering ARIs of 0.3 and 0.5 for the baseline method and Muscle, respectively). **Fig**. S8c-d directly compare the true and the estimated mean methylation matrices and indicate that Muscle’s recovery of the mean methylation matrix better aligns with the true **M** compared to that of the baseline method. In addition, association between the true mean scHi-C ℳ and the Muscle estimate is more evident compared to the estimate from the baseline method (**Fig**. S8e-f).

## 5 Discussion

We presented Muscle as a semi-non negative joint tensor decomposition framework for integrative analysis of chromosome conformation capture and DNA methylation data. Computational experiments with labelled real data and simulated data demonstrated that Muscle’s integrative framework can leverage multiple single cell data modalities to enhance cell type identification. Furthermore, Muscle performed on par or better than existing approaches when presented with single modality scHi-C data. We showcased how Muscle’s parametrization encodes key parameters of interest (cell type specific contact maps, TADs, A/B compartments).

Our application focused on the setting where multiple data modalities are measured simultaneously from individual cells. However, Muscle framework is amenable to extension to cases where individual modalities are separately profiled from cells by employing cell alignment tools such as optimal transport. In applications of Muscle, we observed that the Muscle cell loading parameter that is shared across multiple modalities plays a crucial role in integrative inference. This parameter can be sensitive to the level of variability between the data modalities, necessitating appropriate modeling of the variance terms to balance the contribution of different modalities during integration. We utilized a proportional variance assumption for the scHi-C and methylation modalities and were able to capitalize on the discriminative abilities of the individual modalities for cell types (**Fig**. 3). A more flexible variance modeling approach might be beneficial for integration of additional data modalities. Our implementation of Muscle relied on the Gaussian distribution assumption on the transformed count data. This assumption can be relaxed with tensor models that incorporate Poisson or Negative Binomial distributions (Hong et al., 2020). Another important point in tensor analysis is the selection of the tensor rank. While tensor rank determination is an NP-hard problem compared to matrix rank selection (Håstad, 1990), Fast-Higashi provided a tensor view point on scHi-C without justifying relatively high rank, *R* ≈ 250, in their applications. In Muscle, we employed a heuristic approach to penalize over-fitting; however, a direct regularization term, e.g., group LASSO (Yuan and Lin, 2006), on a collection of rank-1 components could also be employed.

Lastly, while Muscle provides integration, inference, and interpretation advantages compared to alternative methods, its current implementation is relatively slow compared to some of the fast scHi-C analysis methods and warrants further advancement. Specifically, for Lee et al. (2019) scHi-C data at 1Mb resolution, an unoptimized implementation of Muscle required 18 hours (23 cores CPU), while Higashi took 49 hours (10 cores CPU), scHiC Topics took 36 hours (1 core CPU), 3DVI took 4 hours (23 cores GPU), Fast-Higashi took 1 hour (23 cores CPU), scHiCluster took 30 minutes (23 cores CPU), and BandNorm took 15 minutes (23 cores CPU). The speed bottleneck of Muscle is mainly due to additional decomposition steps for estimating loci loadings, which encode key downstream parameters of interests, and warrants further computational developments.

## 6 Acknowledgements

We thank Siqi Shen from the University of Wisconsin-Madison for helping with the raw data processing of scHi-C datasets. We thank both Siqi Shen and Dr. Ye Zheng (Fred Hutchinson Cancer Center) for useful discussions and sharing their analysis. We also thank Dr. Hanbaek Lyu, and Chanwoo Lee, Coleman Breen from the University of Wisconsin-Madison for insightful discussions.

## References

De Lathauwer, L. (2008). Decompositions of a higher-order tensor in block terms—Part II: Definitions and uniqueness. SIAM Journal on Matrix Analysis and Applications 30 (3), 1033–1066.

Gong, Y., C. Lazaris, T. Sakellaropoulos, A. Lozano, P. Kambadur, P. Ntziachristos, I. Aifantis, and A. Tsirigos (2018). Stratification of TAD boundaries reveals preferential insulation of super-enhancers by strong boundaries. Nature communications 9 (1), 1–12.

Håstad, J. (1990). Tensor rank is NP-complete. Journal of Algorithms 11 (4), 644–654.

Hong, D., T. G. Kolda, and J. A. Duersch (2020). Generalized canonical polyadic tensor decomposition. SIAM Review 62 (1), 133–163.

Kiers, H. A., J. M. Ten Berge, and R. Bro (1999). PARAFAC2—Part I. A direct fitting algorithm for the PARAFAC2 model. Journal of Chemometrics: A Journal of the Chemometrics Society 13 (3-4), 275–294.

Kim, H.-J., G. G. Yardımcı, G. Bonora, V. Ramani, J. Liu, R. Qiu, C. Lee, J. Hesson, C. B. Ware, J. Shendure, et al. (2020). Capturing cell type-specific chromatin compartment patterns by applying topic modeling to single-cell Hi-C data. PLoS computational biology 16 (9), e1008173.

Lee, D.-S., C. Luo, J. Zhou, S. Chandran, A. Rivkin, A. Bartlett, J. Nery, C. Fitzpatrick, C. O’Connor, J. Dixon, and J. Ecker (2019, 10). Simultaneous profiling of 3D genome structure and DNA methylation in single human cells. Nature Methods 16, 1–8.

Li, G., Y. Liu, Y. Zhang, N. Kubo, M. Yu, R. Fang, M. Kellis, and B. Ren (2019a, 10). Joint profiling of DNA methylation and chromatin architecture in single cells. Nature Methods 16.

Li, G., Y. Liu, Y. Zhang, N. Kubo, M. Yu, R. Fang, M. Kellis, and B. Ren (2019b). Joint profiling of DNA methylation and chromatin architecture in single cells. Nature methods 16 (10), 991–993.

Li, X., F. Feng, H. Pu, W. Y. Leung, and J. Liu (2021). scHiCTools: A computational toolbox for analyzing single-cell Hi-C data. PLoS computational biology 17 (5), e1008978.

Li, X., G. Zeng, A. Li, and Z. Zhang (2021). DeTOKI identifies and characterizes the dynamics of chromatin TAD-like domains in a single cell. Genome biology 22 (1), 1–26.

Lieberman-Aiden, E., N. L. Van Berkum, L. Williams, M. Imakaev, T. Ragoczy, A. Telling, I. Amit, B. R. Lajoie, P. J. Sabo, M. O. Dorschner, et al. (2009a). Comprehensive mapping of long-range interactions reveals folding principles of the human genome. science 326 (5950), 289–293.

Lieberman-Aiden, E., N. L. Van Berkum, L. Williams, M. Imakaev, T. Ragoczy, A. Telling, I. Amit, B. R. Lajoie, P. J. Sabo, M. O. Dorschner, et al. (2009b). Comprehensive mapping of long-range interactions reveals folding principles of the human genome. science 326 (5950), 289–293.

Lister, R., E. A. Mukamel, J. R. Nery, M. Urich, C. A. Puddifoot, N. D. Johnson, J. Lucero, Y. Huang, A. J. Dwork, M. D. Schultz, et al. (2013). Global epigenomic reconfiguration during mammalian brain development. Science 341 (6146), 1237905.

Liu, H., J. Zhou, W. Tian, C. Luo, A. Bartlett, A. Aldridge, J. Lucero, J. K. Osteen, J. R. Nery, H. Chen, et al. (2021). DNA methylation atlas of the mouse brain at single-cell resolution. Nature 598 (7879), 120–128.

Luo, C., P. Hajkova, and J. R. Ecker (2018). Dynamic DNA methylation: In the right place at the right time. Science 361 (6409), 1336–1340.

Pombo, A. and N. Dillon (2015). Three-dimensional genome architecture: players and mechanisms. Nature reviews Molecular cell biology 16 (4), 245–257.

Ramani, V. and Deng, X., R. Qiu, K. Gunderson, F. Steemers, C. Disteche, W. Noble, Z. Duan, and J. Shendure (2017, 01). Massively multiplex single-cell Hi-C. Nature Methods 14.

Rao, S. S., M. H. Huntley, N. C. Durand, E. K. Stamenova, I. D. Bochkov, J. T. Robinson, A. L. Sanborn, I. Machol, A. D. Omer, E. S. Lander, et al. (2014). A 3D map of the human genome at kilobase resolution reveals principles of chromatin looping. Cell 159 (7), 1665–1680.

Rodriguez, C., J. Borgel, F. Court, G. Cathala, T. Forné, and J. Piette (2010). CTCF is a DNA methylation-sensitive positive regulator of the INK/ARF locus. Biochemical and biophysical research communications 392 (2), 129–134.

Shen, S., Y. Zheng, and S. Keleş (2022). scGAD: single-cell gene associating domain scores for exploratory analysis of scHi-C data. Bioinformatics 38 (14), 3642–3644.

Stevens, T., D. Lando, L. P. Atkinson, Y. Cao, S. Lee, M. Leeb, K. J. Wohlfahrt, W. Boucher, A. O’Shaughnessy-Kirwan, J. Cramard, A. J. Faure, M. Ralser, E. Blanco, L. Morey, M. Sanso, M. G. S. Palayret, B. Lehner, L. Di Croce, and E. Laue (2017, 03). 3D structure of individual mammalian genomes studied by single cell Hi-C. Nature 544, 59–64.

Tan, L., W. Ma, H. Wu, Y. Zheng, D. Xing, R. Chen, X. Li, N. Daley, K. Deisseroth, and X. S. Xie (2021). Changes in genome architecture and transcriptional dynamics progress independently of sensory experience during post-natal brain development. Cell 184 (3), 741–758.

Ulianov, S. V., V. V. Zakharova, A. A. Galitsyna, P. I. Kos, K. E. Polovnikov, I. M. Flyamer, E. A. Mikhaleva, E. E. Khrameeva, D. Germini, M. D. Logacheva, et al. (2021). Order and stochasticity in the folding of individual Drosophila genomes. Nature communications 12 (1), 1–17.

Wang, H., M. T. Maurano, H. Qu, K. E. Varley, J. Gertz, F. Pauli, K. Lee, T. Canfield, M. Weaver, R. Sandstrom, et al. (2012). Widespread plasticity in CTCF occupancy linked to DNA methylation. Genome research 22 (9), 1680–1688.

Yang, T., F. Zhang, G. G. Yardımcı, F. Song, R. C. Hardison, W. S. Noble, F. Yue, and Q. Li (2017). HiCRep: assessing the reproducibility of Hi-C data using a stratum-adjusted correlation coefficient. Genome research 27 (11), 1939–1949.

Yu, M., A. Abnousi, Y. Zhang, G. Li, L. Lee, Z. Chen, R. Fang, T. M. Lagler, Y. Yang, J. Wen, et al. (2021). Snaphic: A computational pipeline to identify chromatin loops from single-cell hi-c data. Nature methods 18 (9), 1056–1059.

Yu, M. and B. Ren (2017). The three-dimensional organization of mammalian genomes. Annual review of cell and developmental biology 33, 265.

Yuan, M. and Y. Lin (2006). Model selection and estimation in regression with grouped variables. Journal of the Royal Statistical Society: Series B (Statistical Methodology) 68 (1), 49–67.

Zhang, R., T. Zhou, and J. Ma (2022a). Multiscale and integrative single-cell Hi-C analysis with Higashi. Nature biotechnology 40 (2), 254–261.

Zhang, R., T. Zhou, and J. Ma (2022b). Ultrafast and interpretable single-cell 3D genome analysis with Fast-Higashi. In International Conference on Research in Computational Molecular Biology, pp. 300–301. Springer.

Zheng, Y., S. Shen, and S. Keleş (2022). Normalization and de-noising of single-cell hi-c data with bandnorm and scvi-3d. Genome Biology 23 (1), 1–34.

Zhou, J., J. Ma, Y. Chen, C. Cheng, B. Bao, J. Peng, T. J. Sejnowski, J. R. Dixon, and J. R. Ecker (2019). Robust single-cell Hi-C clustering by convolution-and random-walk–based imputation. Proceedings of the National Academy of Sciences 116 (28), 14011–14018.

